# 4D Biomimetic Morphing Hydrogel Scaffold via Biaxial Gradient Programming

**DOI:** 10.64898/2026.07.28.741270

**Authors:** Aixiang Ding, Kaelyn L. Gasvoda, Eben Alsberg

**Affiliations:** Richard and Loan Hill Department of Biomedical Engineering, University of Illinois at Chicago, 909 S. Wolcott Avenue, Chicago, IL 60612, USA. Emails: (A.D.); (E.A.); Jesse Brown Veterans Affairs Medical Center (JBVAMC), Chicago, IL 60612, USA; Departments of Mechanical & Industrial Engineering, Orthopaedic Surgery, and Pharmacology and Regenerative Medicine, University of Illinois at Chicago, 909 S. Wolcott Avenue, Chicago, IL 60612, USA

**Keywords:** 4D printing, biofabrication, hydrogel, functional gradient materials, tissue engineering

## Abstract

Four-dimensional (4D) materials incorporating functional gradient designs offer a powerful platform for engineering dynamic structures capable of programmed shape transformations in response to environmental stimuli. However, most gradient-based 4D systems rely on uniaxial gradients, which typically generate simple, symmetric deformations with uniform curvature, limiting their ability to recreate biomimetic architectures that require spatially coordinated morphogenesis. Here, we report a biaxial gradient-engineered 4D hydrogel system capable of programmable, non-uniform shape morphing within a single construct. A one-step photocrosslinking strategy integrates vertical light attenuation and horizontal grayscale photomask patterning to establish orthogonal crosslinking gradients along two directions, producing spatially heterogeneous swelling stresses that drive controlled multi-directional deformation. The resulting hydrogels exhibit tunable swelling and mechanical properties, enabling precise regulation of curvature distribution and shape transformation. This biaxial gradient platform generates diverse biomimetic architectures, including swan-neck, fiddlehead fern, sea star, and *Euonymus europaeus*-like structures. Importantly, the system supports cell-laden biofabrication, where human mesenchymal stem cell-encapsulated constructs maintain high viability and undergo chondrogenic differentiation while preserving programmed morphologies. This work establishes biaxial gradient-programmed 4D hydrogels as a robust strategy for integrating morphogenesis with tissue formation, advancing biomimetic biofabrication and morphogenetic tissue engineering.

## Introduction

Four-dimensional (4D) materials are engineered systems that can be capable of undergoing programmed shape transformations in response to external stimuli, such as temperature, moisture, pH, light, or electric/magnetic fields, or internal stimuli, including cell-generated contractile forces when living cells are seeded with the constructs^1–4^. These dynamic materials can be fabricated from shape memory polymers, hydrogels, liquid crystalline elastomers, and their composites through manufacturing approaches such as additive printing, casting, molding, and modular assembly^5–7^. Owing to their ability to transition from static to adaptive architectures, 4D materials have attracted significant attention across soft robotics, deployable devices, biomedical engineering, and bioinspired systems^8–10^.

In tissue engineering, 4D materials are increasingly recognized as a next-generation platform because they enable the generation of complex, dynamically evolving architectures that are otherwise difficult to achieve using conventional static scaffolds^11–14^. Beyond structural reconfiguration, dynamically morphing systems can enhance tissue regeneration and guide cellular distribution by providing temporally regulated mechanical and geometric cues^15, 16^.

Importantly, such systems offer a powerful experimental framework for recapitulating key aspects of tissue morphogenesis, thereby enabling the study of physiological developmental processes *in vitro*^17, 18^.

Strategies for constructing 4D materials typically rely on multilayer assemblies with spatially mismatched stress-strain distributions^19, 20^ or composite systems incorporating microstructural anisotropy, such as aligned micro- or nanofiber reinforcements^21–23^. While these approaches can achieve predictable bending, twisting, or folding behaviors, they often involve structural complexity and may suffer from practical limitations, including interfacial delamination in layered systems or limited control over fiber distribution and alignment.

More recently, functional gradient materials, characterized by spatial variations in crosslinking density, composition, or mechanical properties, have emerged as an elegant and structurally simple strategy for 4D design^24–26^. Gradients can be generated through techniques such as attenuated photopolymerization, controlled diffusion or phase separation, and microfluidic processing^27–30^. These systems allow tunable gradient sharpness and amplitude, enabling programmable deformation magnitudes and improved fabrication versatility. Moreover, the resulting spatial mechanical heterogeneity may provide biophysical cues potentially capable of modulating cell behavior and further enhancing the constructs’ relevance for regenerative medicine applications.

Despite these advances, most existing gradient-based 4D systems rely on uniaxial gradients^31–33^. Consequently, their morphing behavior is typically restricted to simple, symmetric deformations that generate uniform curvature distributions and geometrically elementary forms. For example, a single-layer hydrogel strip containing a through-thickness crosslinking gradient commonly undergoes monomodal bending upon swelling, yielding a simple “C”-shaped configuration (**Figure 1a**). Although effective for demonstrating proof-of-concept actuation, such uniform curvature limits the ability to recreate biomimetic architectures, which often require coordinated, spatially non-uniform curvature distributions and multi-regional deformation patterns to emulate natural morphogenesis^34^. In biological tissues, curvature is rarely homogeneous; instead, it varies spatially to generate hierarchical forms and region-specific biochemical and mechanical environments. These features are challenging to be captured by single-axis gradient systems.

**Figure 1.**
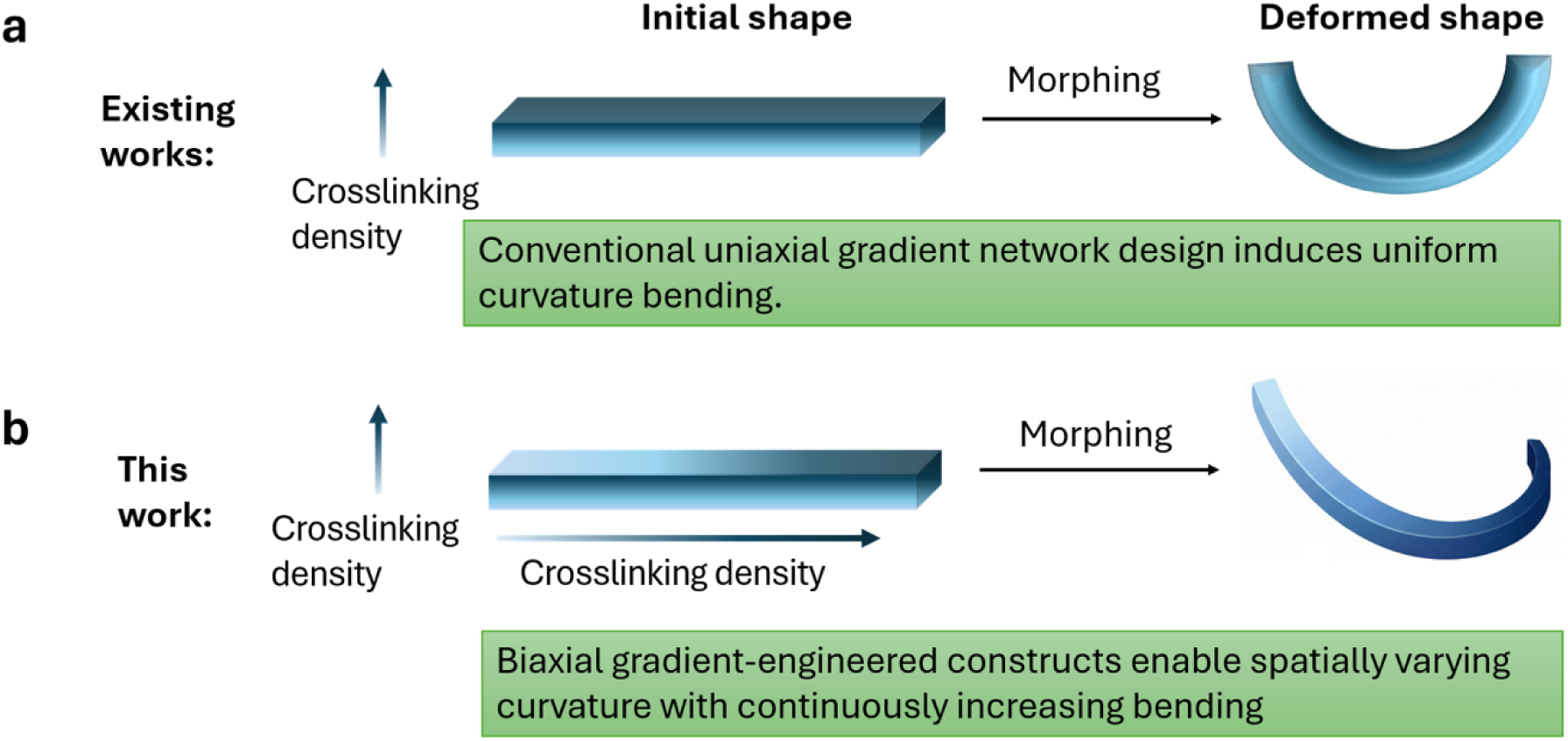
Schematic comparison between (a) a conventional uniaxial gradient hydrogel system that induces uniform-curvature bending upon stimulation and (b) the engineered biaxial gradient system that enables spatially coordinated, non-uniform curvature formation within a single construct.

To overcome these limitations, a 4D hydrogel system capable of programmable, non-uniform deformation within a single construct to generate spatially varying curvatures and architecturally sophisticated forms has been developed. Specifically, a biaxial gradient configuration that introduces controlled gradient crosslinking density along both the horizontal (x-axis) and vertical (z-axis) directions within a single hydrogel construct was engineered (**Figure 1b**). This orthogonal gradient architecture generates spatially heterogeneous swelling stresses in multiple directions, thereby enabling coordinated, non-uniform shape transformations across the construct. Using this platform, the formation of complex curvature architectures with high spatial precision and reproducibility were demonstrated. Furthermore, by encapsulating human mesenchymal stem cells (hMSCs) within these dynamically reconfigurable matrices, the feasibility of coupling spatially programmed multi-axial morphogenesis with lineage-specific tissue development was illustrated.

## Results and Discussion

### Bioinks synthesis and characterization

Oxidized and methacrylated alginate (OMA)^35^ was synthesized and used as the primary polymer for hydrogel fabrication (**Figure S1**). Calcium ion (Ca^2+^)-crosslinked OMA hydrogels were subsequently processed into a jammed microgel slurry (**Figure S3a,b**) using our previously established mechanical fragmentation approach^36, 37^. Owing to the presence of free methacrylate groups, the slurry microgels can be further photocrosslinked in the presence of a photoinitiator to form stable bulk hydrogels (**Figure S3c,d**). Although OMA provides excellent cytocompatibility as a scaffold material, alginate lacks intrinsic cell-adhesive motifs that are critical for promoting cell-matrix interactions^38, 39^. To address this limitation, methacrylated gelatin (GelMA, **Figure S2**) was incorporated into the OMA slurry microgels to formulate the final bioink. GelMA contains abundant RGD peptide motifs, enabling cell adhesion and enhancing cellular interaction with the hydrogel matrix^40^.

The resulting OMA/GelMA bioink exhibited favorable rheological properties that facilitate extrusion-based printing of freestanding constructs. Specifically, the bioink displayed a storage modulus (G′) consistently higher than the loss modulus (G″) across the tested frequency range (1% 10%), indicating dominant solid-like behavior at low oscillatory frequencies (**Figure 2a**). A pronounced shear-thinning behavior critical for extrusion printing was evidenced by the crossover of G′ and G″ at approximately 30% shear strain (**Figure 2b**), indicating a transition from solid-like to liquid-like behavior under applied strain. Furthermore, the viscosity decreased with increasing shear rate (**Figure 2c**), further confirming the shear-thinning characteristics that enable smooth extrusion during the printing process.

**Figure 2.**
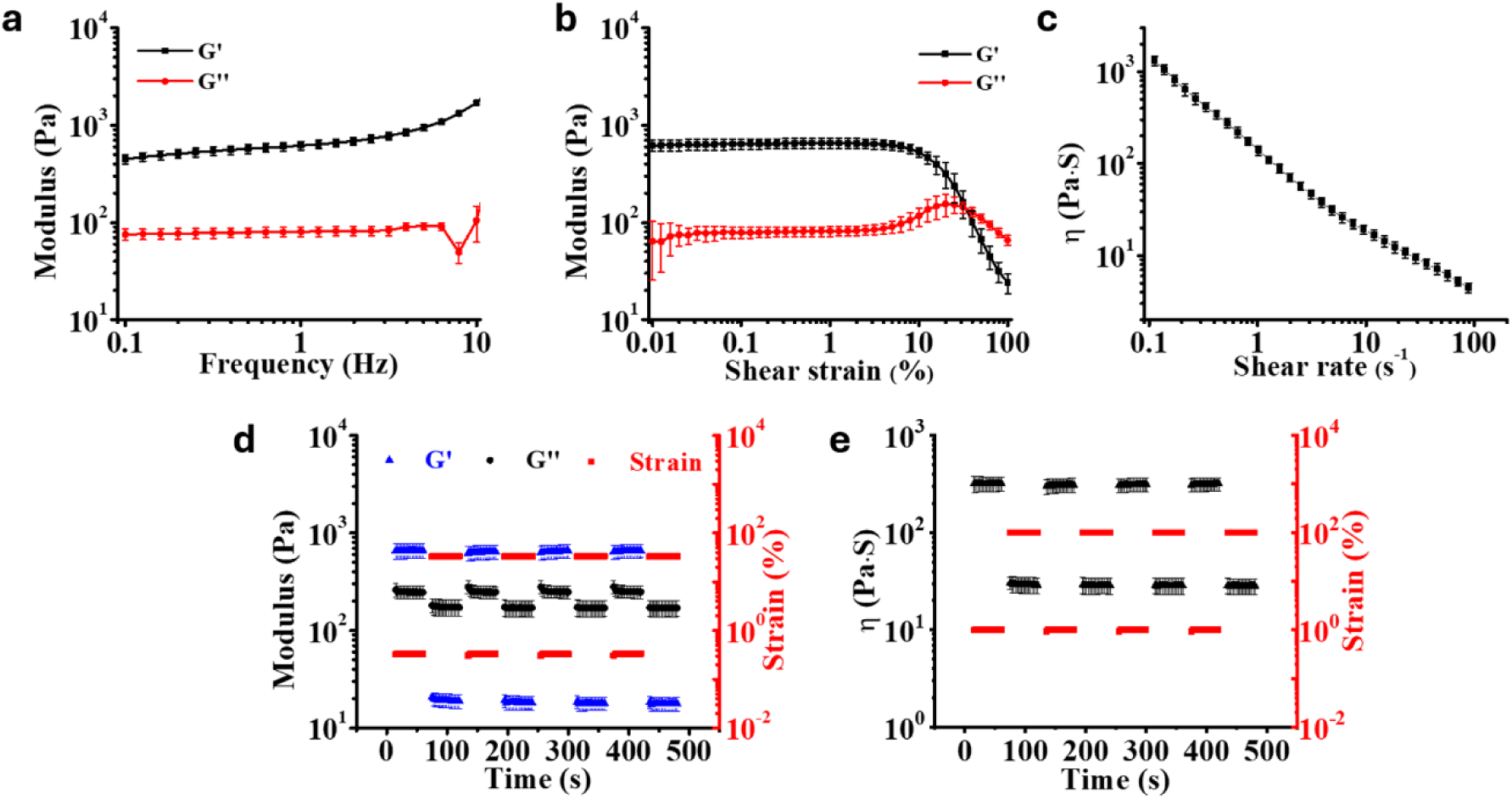
Rheological characterization of the OMA/GelMA bioinks. (a) Storage modulus (G′) and loss modulus (G″) of the bioink as a function of oscillatory frequency. Shear-thinning behavior illustrated by the (b) crossover of G′ and G″ with increasing shear strain and (c) decrease in viscosity with increasing shear rate. Reversible transitions of (d) G′ and G″ and (e) viscosity under cyclic strain switching between 1% and 100%. Data are presented as mean ± standard deviation (±SD), *N* = 3.

Self-healing behavior is another critical property for maintaining structural stability after extrusion. To evaluate this property, the modulus and viscosity of the bioink were measured under cyclic oscillatory strains alternating between 1% and 100% strain. The bioink demonstrated rapid and reversible transitions in both modulus and viscosity. Specifically, the material exhibited solid-like behavior (G′ > G″) and high viscosity (∼1000 Pa·s) at 1% strain, while showing liquid-like behavior (G′ < G″) and low viscosity (∼10 Pa·s) at 100% strain (**Figure 2d,e**). Importantly, these transitions occurred rapidly and reproducibly over at least five consecutive cycles without noticeable fatigue, demonstrating excellent self-healing capability.

### Printability assessment

Benefiting from these favorable rheological properties, the OMA/GelMA bioink demonstrated excellent printability. For example, extrusion through a 22-gauge needle (inner diameter 0.41 mm) produced a stable filament suspended in air without collapse, indicating strong filament-forming capacity (**Figure 3a**). Under optimized printing parameters (**Table S1**), centimeter-scale freestanding constructs were successfully printed, including a cube, a hollow cylinder, and a grid structure (**Figure 3b, Movie S1**). The printed constructs maintained their structural integrity immediately after deposition and could be further stabilized through photocrosslinking, resulting in structures with high fidelity (**Figure 3c,d**). Notably, slight distortion was observed at the corners of the printed grid after photocrosslinking, which resulted from transferring the construct from the printing platform to a quartz plate prior to UV exposure rather than from printing instability.

**Figure 3.**
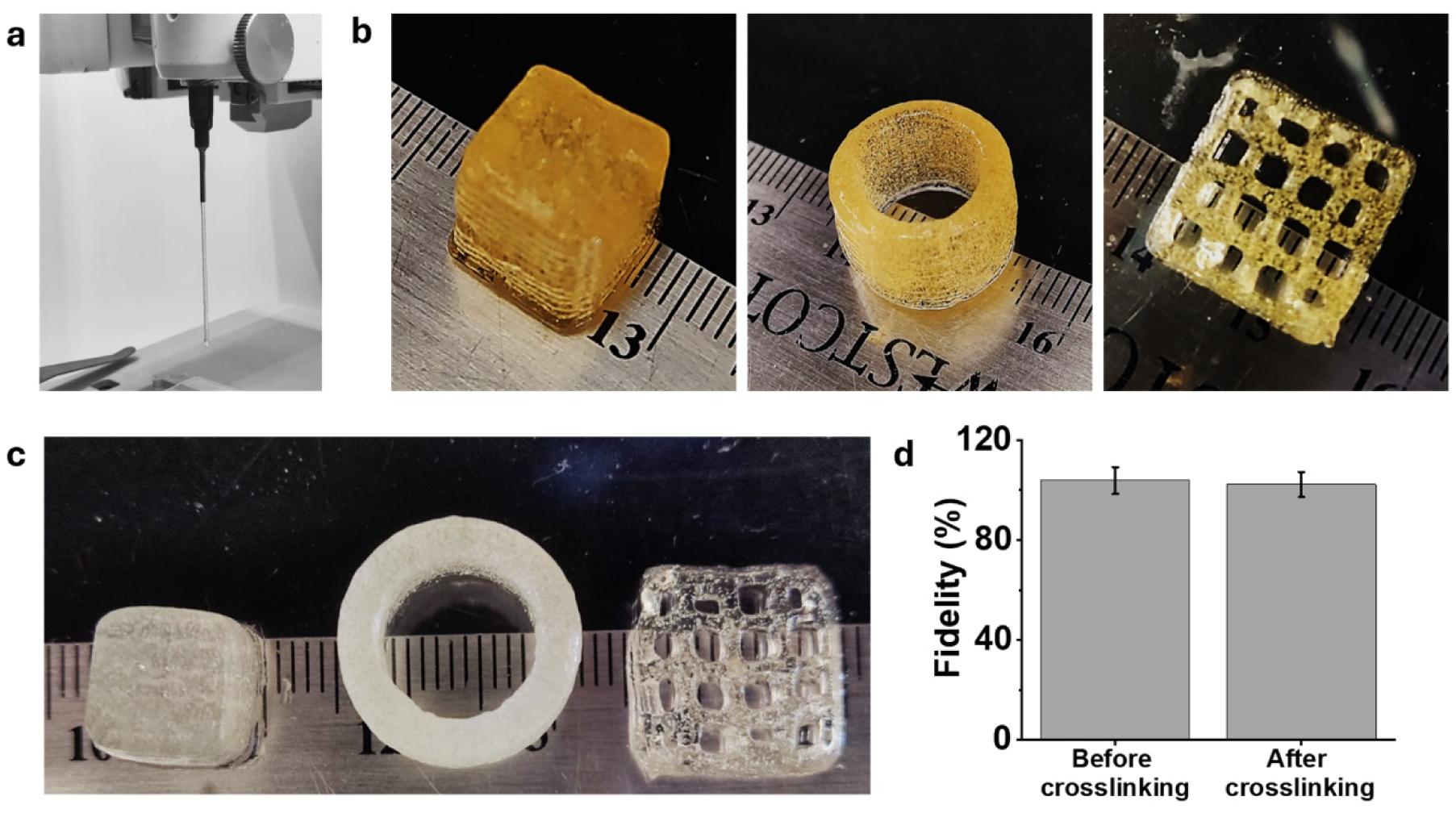
Printability evaluation of the OMA/GelMA bioink. (a) Photograph of a stable filament extruded from a 22-gauge needle suspended in air. Images of printed 3D constructs, including a cube (10 × 10 × 10 mm), hollow cylinder (outer diameter 15 mm, inner diameter 11.6 mm, height 10 mm), and grid (20 mm × 20 mm × 4 mm), (b) before and (c) after photocrosslinking (20 mW cm-2, 30 s). (d) Printing fidelity (%) of the cube structure before and after photocrosslinking.

### Biaxial gradient engineering

To generate a biaxial gradient, a one-step photocrosslinking strategy was developed for the printed constructs (**Figure 4a**). Using a representative strip geometry as an example, and following our previously established approach^32, 36, 41^, a photoabsorber (PA) was incorporated into the printed precursor to attenuate light penetration along the vertical (z) direction, thereby producing a through-thickness crosslinking gradient, referred to as the vertical gradient (VG).

**Figure 4.**
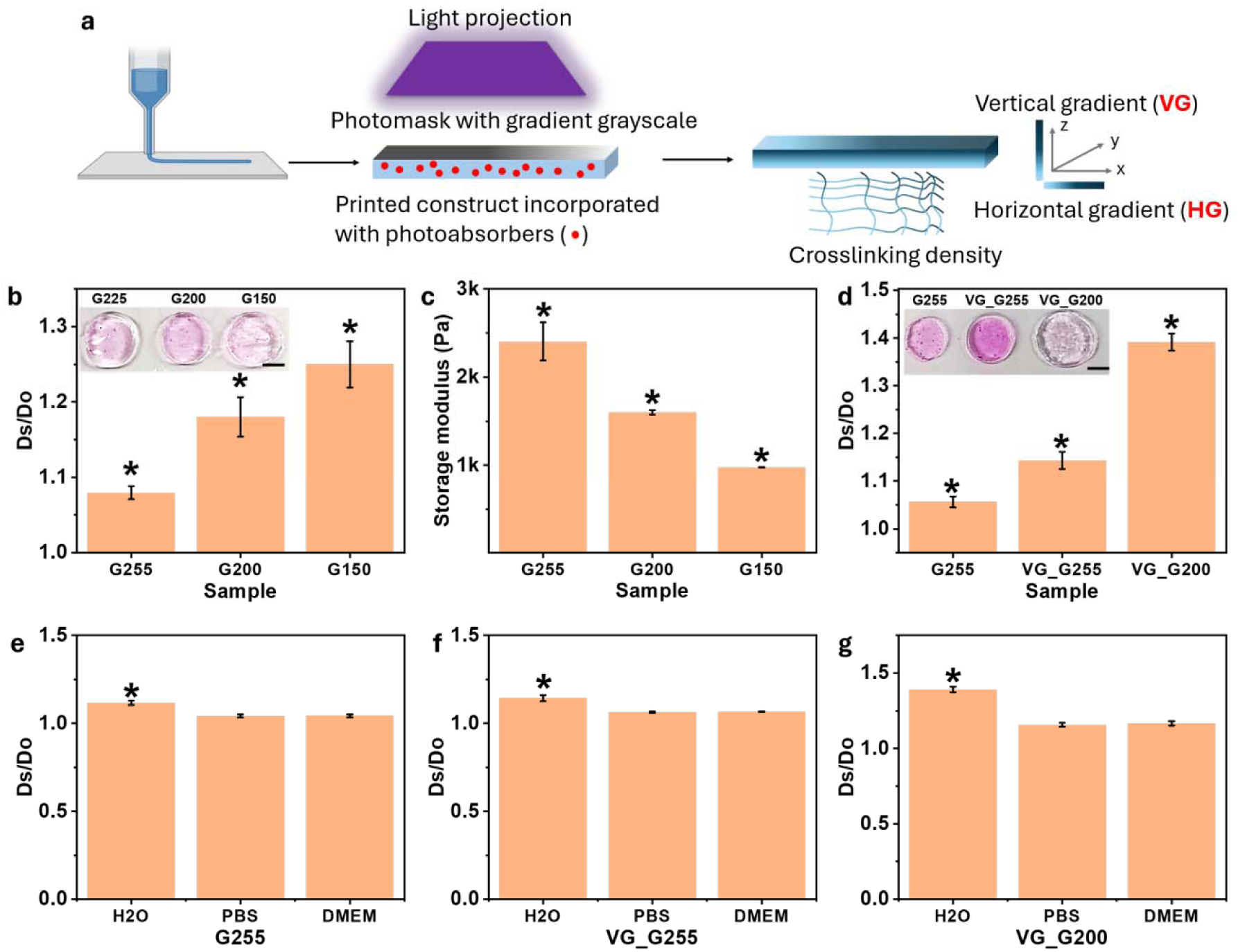
Biaxial gradient engineering. (a) Schematic illustration of biaxial gradient formation via a one-step photocrosslinking strategy that combines vertical light attenuation and horizontal grayscale masking. (b) Volumetric swelling ratios and (c) storage modulus of hydrogels photocrosslinked under different uniform grayscale photomasks without PA incorporation (no VG formation) after swelling in H_2_O. (d) Volumetric swelling ratios of hydrogels photocrosslinked under different uniform grayscale photomasks with PA incorporation (VG formation) after swelling in H_2_O. (e–f) Swelling behavior of hydrogels photocrosslinked under uniform G255 photomasks (e) without and (f) with VG formation after incubation in H_2_O, PBS, and DMEM. (g) Swelling behavior of hydrogels photocrosslinked under uniform G200 photomasks with VG formation after incubation in H2O, PBS, and DMEM. Insets in (b) and (d) show representative images of disc-shaped hydrogels after swelling in H2O. Hydrogels were supplemented with 0.005% w/v rhodamine B to aid visualization. Ds denotes the diameter of the swollen hydrogel disc, and Do denotes the initial diameter (8 mm). \**p* < 0.05 for (b–c); \**p* < 0.05 compared with unlabeled groups in (e–g). Scale bars = 5 mm. Data are presented as mean ± standard deviation (±SD), *N* = 3.

Simultaneously, a photomask with a graded grayscale pattern was positioned above the precursor solution to introduce spatial variations in light transmittance along the horizontal (x) direction, generating a horizontal gradient (HG). Through the combined effects of z-direction light attenuation and x-direction masking, a biaxial crosslinking gradient can be established within a single hydrogel construct.

For HG engineering, photomasks with grayscale values ranging from 0 to 255 (G0– G255) were used to regulate light transmittance. A grayscale value of 255 (G255) corresponds to full transparency with 100% light transmission, whereas G0 represents complete opacity with full light blocking. Therefore, tuning the grayscale value range of the photomask provides a straightforward means to modulate the range of the horizontal crosslinking gradient.

To evaluate the effect of photomask grayscale on hydrogel swelling and mechanical properties, hydrogels were first synthesized using photomasks with uniform grayscale values of G255, G200, G150, G100, and G50 (**Figure S4**) without incorporating the PA (i.e., no VG formation).

Hydrogels prepared under G50 failed to maintain structural integrity after photocrosslinking (**Figure S5a**). Although hydrogels could be formed under G100, they were mechanically fragile and prone to disintegration when transferred to tissue culture plates. Therefore, hydrogels formed under G100 and G50 conditions were excluded from further investigation.

Hydrogels prepared with G255, G200, and G150 photomasks exhibited progressively increased volumetric swelling ratios with decreasing grayscale values (**Figure 4b**). This behavior can be attributed to reduced crosslinking density resulting from decreased light transmission through the photomask. Correspondingly, these hydrogels displayed a significant decrease in storage modulus with decreasing grayscale values (**Figure 4c**). These results confirm that grayscale modulation can effectively regulate crosslinking density, suggesting that a continuous HG in network structure can be generated using a photomask with spatially varying grayscale values.

Next, the influence of incorporating a VG on hydrogel swelling and mechanical properties was investigated for constructs fabricated under G255 and G200 conditions (VG_G255 and VG_G200). The VG_G150 condition was not evaluated because stable hydrogels could not be formed under this condition (**Figure S5b**). Incorporation of a VG significantly increased hydrogel swelling (**Figure 4d**). For example, the volumetric expansion of VG_G255 increased to 1.14, compared with 1.05 for G255 without VG. The swelling ratio further increased to 1.39 when the grayscale value was reduced to G200 (VG_G200). Correspondingly, hydrogels incorporating VG also exhibited a significant reduction in storage modulus (**Figure S6**), consistent with the formation of an overall lower crosslinking density within the gradient networks.

In addition, the hydrogels exhibited different swelling behaviors in various aqueous environments, including deionized water (H_2_O), phosphate-buffered saline (PBS, pH 7.4), and Dulbecco’s Modified Eagle’s Medium (DMEM) (**Figure 4e–g**). Regardless of the presence of VG, all hydrogels showed maximum swelling in H_2_O and comparatively lower but similar swelling in PBS and DMEM, which can be attributed to the ionic strength of the buffered media that screens electrostatic repulsion between alginate chains and suppresses hydrogel expansion^42^.

### Shape morphing evaluation

To evaluate the shape-morphing capability of the constructs, strip-shaped hydrogels with different biaxial crosslinking gradient configurations were fabricated and cultured in PBS at 37 °C for 12 h, after which images were collected for analysis. For comparison, hydrogel strips containing single gradients, either HG or VG crosslinking, were fabricated and cultured under the same conditions.

Hydrogel strips presenting only HG crosslinking were fabricated using gradient grayscale photomasks without incorporating PAs in the slurry bioinks. These strips exhibited minimal shape change and largely maintained a near-straight configuration after incubation (**Figure 5a**), suggesting that the HG alone did not induce significant shape morphing under these conditions.

**Figure 5.**
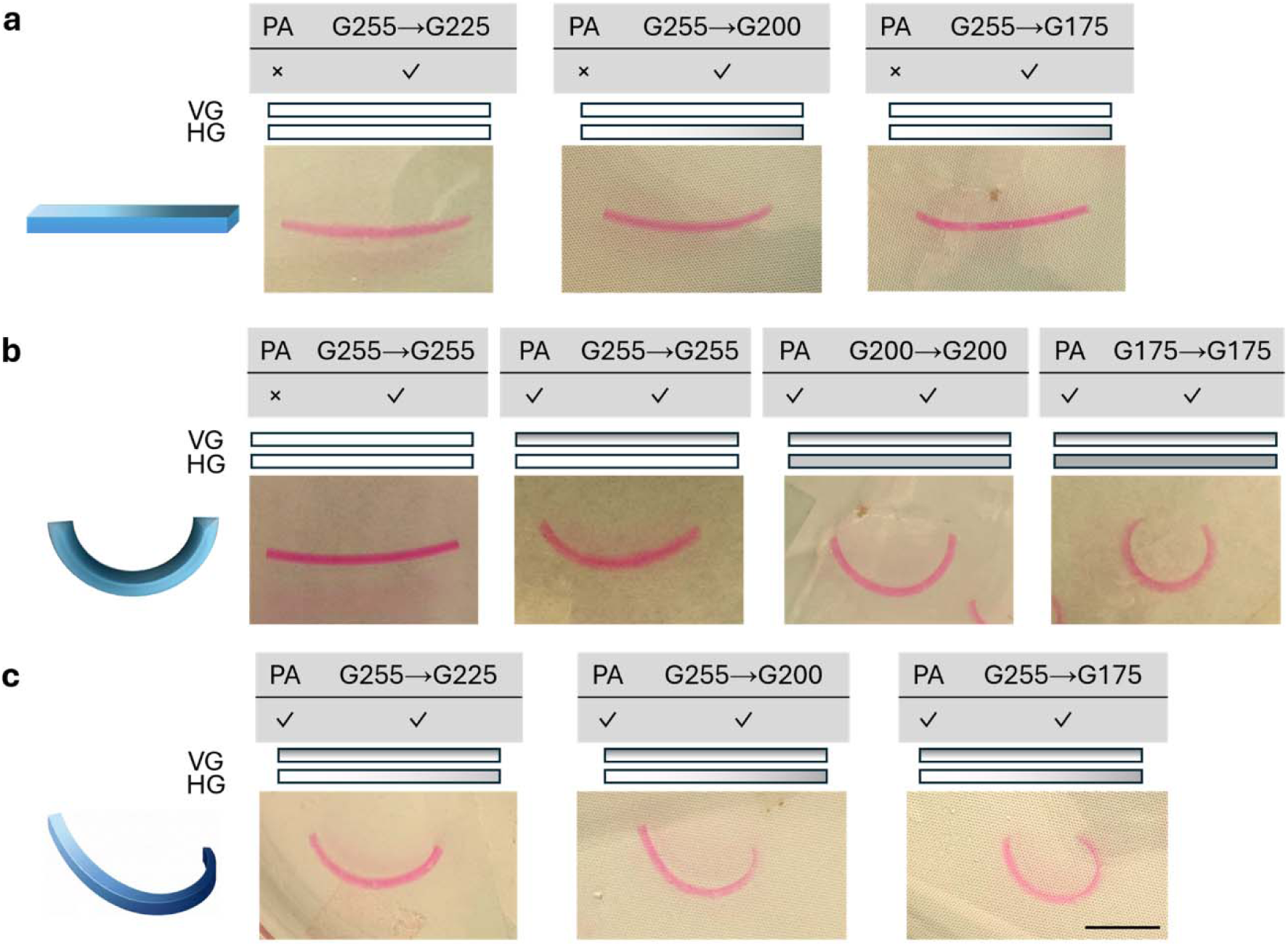
Shape morphing of strip hydrogels with different gradient configurations. Representative images of hydrogel strips exhibiting (a) HG only, (b) VG only, and (c) VG_HG biaxial gradients. Scale bar = 10 mm.

Hydrogel strips presenting only VG crosslinking were fabricated using homogeneous grayscale photomasks together with PA incorporation. These strips exhibited uniform bending, forming typical “C”-shaped configurations (**Figure 5b**). Moreover, the bending curvature increased with decreasing photomask grayscale values. This behavior could be attributed to the reduced light intensity transmitted through darker photomasks, which decreases crosslinking density and enhances hydrogel swelling, thereby generating larger swelling-induced strain mismatches across construct thickness. These observations are consistent with previously reported gradient hydrogel systems^32, 41^.

In contrast, hydrogel strips incorporating orthogonal biaxial gradients (VG_HG) exhibited non-uniform bending, forming curved structures with spatially varying curvature along the strip (**Figure 5c**). The overall curvature increased as the HG range increased, indicating that the horizontal gradient effectively modulates the spatial distribution of swelling-induced stresses. Notably, the curvature progressively increased toward regions corresponding to lower grayscale values, where crosslinking density is lower and swelling is greater. For example, in the VG_G255→G225 configuration, the largest curvature was observed near the G225 end of the strip. These results demonstrate that biaxial gradient engineering enabled spatially coordinated deformation within a single hydrogel construct, allowing the generation of non-uniform curvature distributions that cannot be achieved with single-gradient systems.

### Complex architectural engineering via multi-directional morphing

The biaxial gradient system was next applied to engineer more complex architectures through spatially coordinated deformations. Using strip-shaped hydrogels as a model system, a constant VG was maintained while introducing two distinct HG orientations: one with the grayscale transitioning from transparent to opaque toward the strip center, and the other with the opposite transition from opaque to transparent toward both ends. This design enabled the formation of multiple, discontinuous curvatures within a single construct (**Figure 6a**).

**Figure 6.**
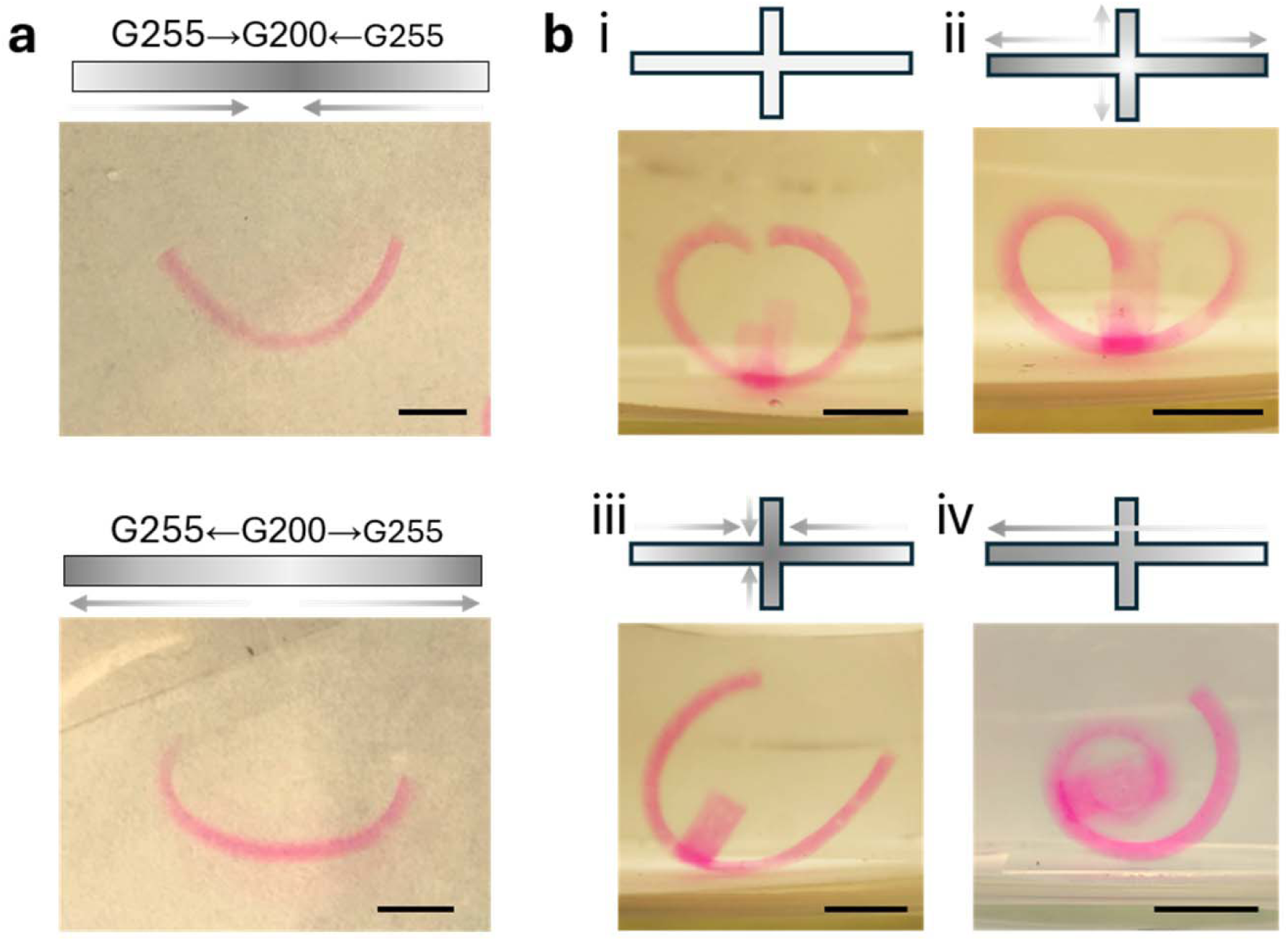
Complex architecture formation of biaxial gradient hydrogels with a VG and different HG configurations. (a) Shape morphing of strip hydrogels incorporating two opposite HG orientations. (b) Shape morphing of cross-shaped hydrogels with different HG configurations in combination with a VG. Scale bars = 5 mm.

Specifically, when the two HGs were configured with grayscale transitioning from transparent at the strip ends to opaque toward the center, both ends of the strip exhibited greater curvature due to the relatively lower crosslinking density in those regions. In contrast, when the gradients configured with the opposite grayscale transition, a more pronounced curvature developed at the center of the construct. These results demonstrate that the direction of horizontal gradients could effectively modulate the distribution of bending curvature along the hydrogel strip.

This concept was further extended by altering the initial geometry from a simple strip to a cross-shaped hydrogel construct. By spatially defining the gradient crosslinking orientations within the x–y plane, a variety of complex architectures were generated through coordinated deformation (**Figure 6b,S7**). For example, a cross-shaped construct incorporating only VG crosslinking produced a baseline morphology with all arms curling upward after swelling (**Figure 6b, i**). When two HGs were introduced in opposite directions along the cross arms, the resulting structures displayed distinct curvature formation in each arm (**Figure 6b, ii and iii**). Specifically, greater curvature was observed at the arm ends when increased light blockage was applied toward the ends (**Figure 6b, ii**), whereas reduced end curvature was observed when the opposite HG configuration was applied (**Figure 6b, iii**). Conversely, introducing a single HG oriented toward one side generated a different asymmetric architecture (**Figure 6b, iv**). It should be noted that differences in the bending behavior of the vertical arms were not readily visible due to their substantially shorter lengths relative to the horizontal arms, as well as limitations associated with the viewing angle. These results highlight the versatility of biaxial gradient engineering in programming spatially complex morphing behaviors, enabling the generation of architecturally sophisticated hydrogel structures through controlled multi-directional deformation.

### Biomimetic 4D fabrication

In nature, many biological structures arise through spatially coordinated morphogenesis, where heterogeneous mechanical stresses orchestrate the development of complex 3D geometries from relatively simple initial forms^43^. Replicating such morphogenetic processes has become a central objective in biomimetic engineering using 4D materials, as programmable shape morphing provides a powerful route to transform initially planar or simple constructs into architecturally sophisticated structures^44, 45^. Building on the capability of the biaxial gradient strategy to generate spatially non-uniform bending and multi-directional morphing, this platform was further extended toward biomimetic 4D fabrication. Using this approach, a series of structures inspired by natural morphologies were successfully generated, including a swan-neck shape, a fiddlehead fern, a sea star, and a *Euonymus europaeus* flower (**Figure 7**). These biomimetic architectures emerge from the programmable spatial distribution of crosslinking gradients (**Figure S8**), which produces coordinated deformation patterns that emulate natural shape formation processes. The ability to engineer such biomimetic morphologies highlights the versatility of the biaxial gradient system for recreating complex biological architectures.

**Figure 7.**
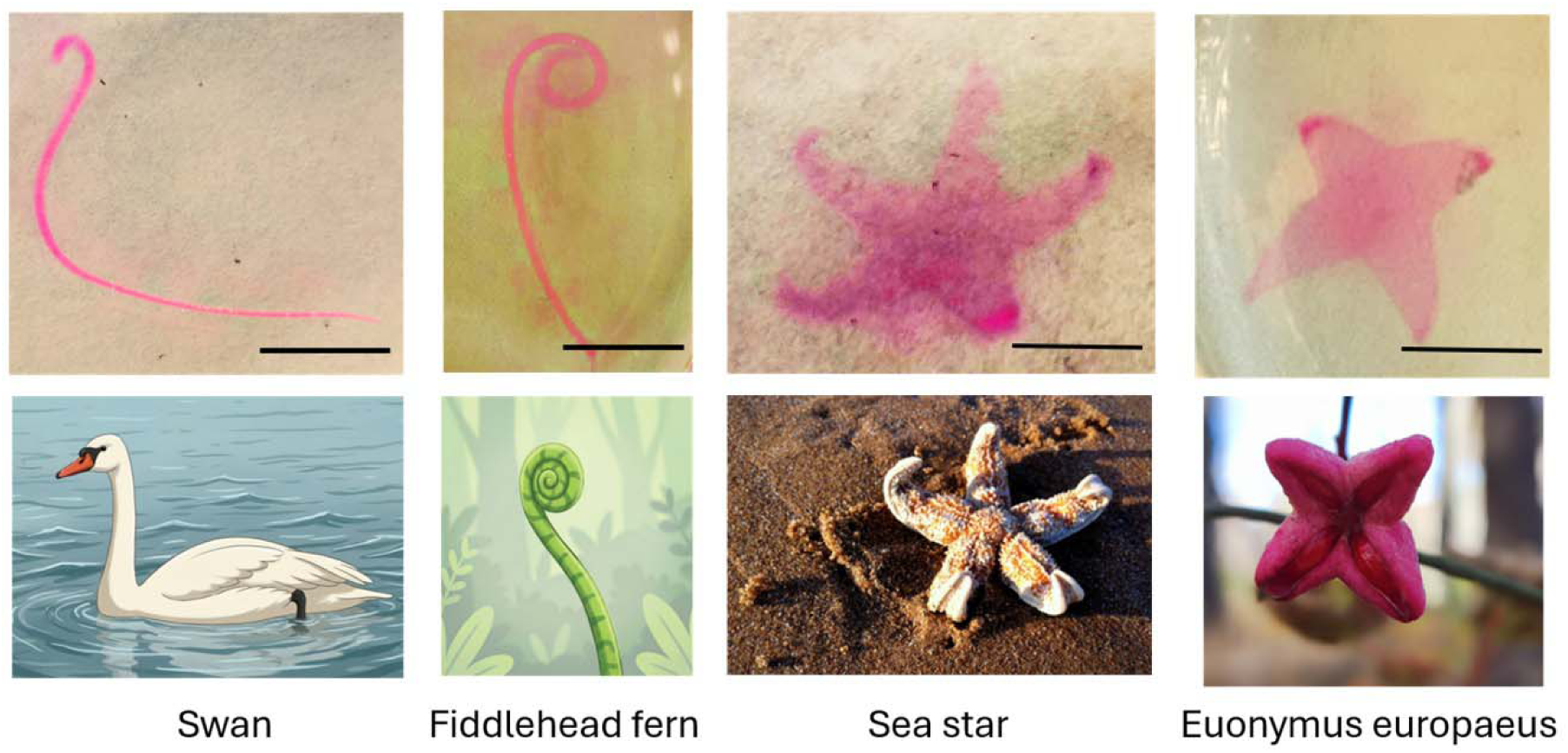
Biomimetic 4D architectures generated using the biaxial gradient hydrogel system. Representative examples of biomimetic structures, including a swan-neck shape, fiddlehead fern, sea star, and *Euonymus europaeus* flower formed through programmed multi-directional morphing. The sea star image is credited to Jane Dallaway. Scale bars = 10 mm.

### 4D morphogenetic tissue engineering

Morphogenetic tissue engineering represents an important paradigm in regenerative medicine, aiming to recapitulate the developmental processes through which native tissues form via coordinated shape morphogenesis, cell differentiation, and extracellular matrix (ECM) deposition and remodeling^46, 47^. 4D biomaterials provide a promising platform to integrate programmable structural morphing with biological tissue formation, enabling dynamic scaffolds that guide both tissue geometry and cellular development over time.

Finally, this platform was applied to morphogenetic tissue engineering, demonstrated through a proof-of-concept study of 4D cartilage-like tissue formation. hMSCs were encapsulated within the hydrogel strips, which were then processed under different gradient configurations and divided into four groups: Ctrl1, Ctrl2, Ctrl3, and Exp. Ctrl1 served as the control group without either VG or HG. Ctrl2 represented the group presenting only HG, while Ctrl3 represented the group presenting only VG. The Exp group corresponded to a biaxial gradient configuration (VG_HG). All constructs were subsequently cultured in chondrogenic differentiation medium for 21 days.

As expected, Ctrl1 and Ctrl2 strips exhibited minimal shape changes throughout the culture period, whereas Ctrl3 strips underwent uniform bending, forming typical C-shaped configurations. In contrast, Exp strips displayed non-uniform morphing, forming curved structures with continuously increasing curvature toward the lower grayscale end (**Figure 8a**). All constructs maintained excellent structural stability throughout the 21-day culture period, with no noticeable structural degradation. Moreover, the encapsulated cells exhibited a predominantly round morphology (**Figure S9**) and remained highly viable, as indicated by the predominance of green fluorescence in the live/dead assay-stained images (**Figure 8b**), demonstrating the excellent cytocompatibility of the dynamic hydrogel system.

**Figure 8.**
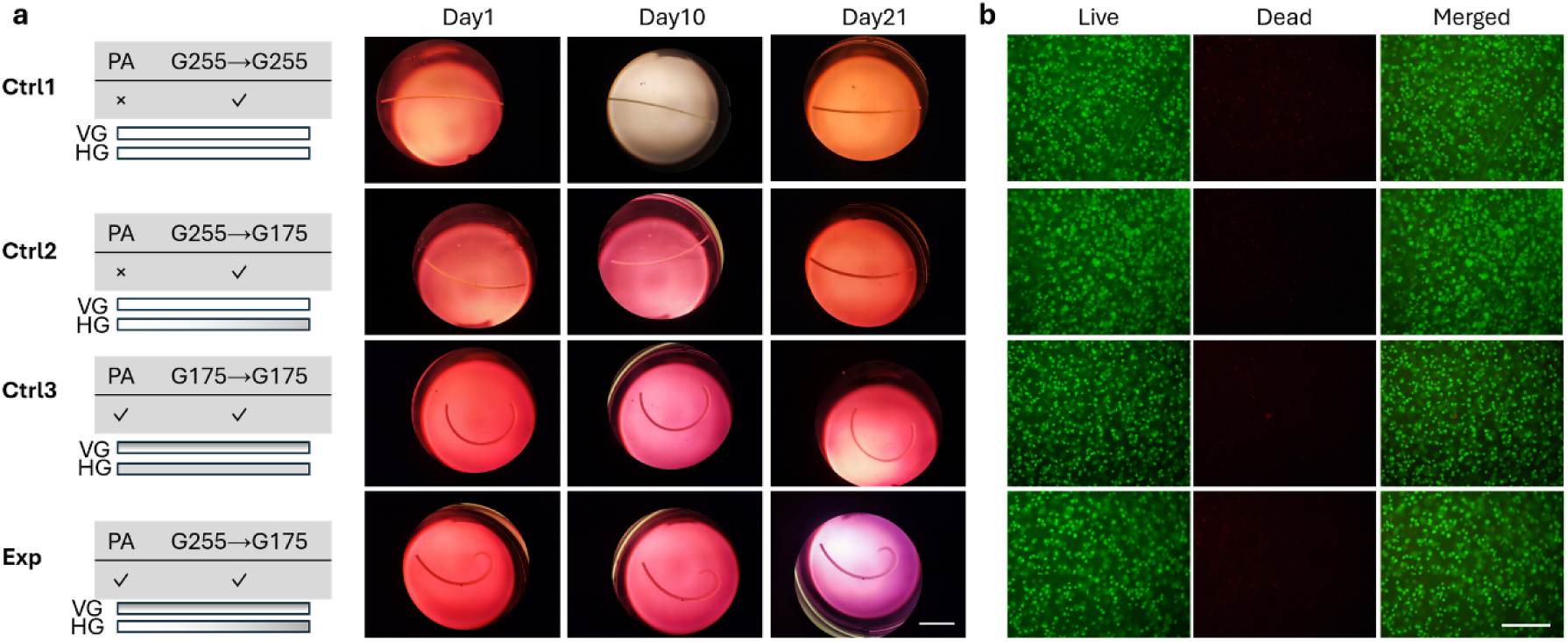
Shape morphing and cell viability of cell-laden constructs during chondrogenic culture. (a) Representative images of cell-laden hydrogel strips from different groups at Day 1, Day 10, and Day 21 during culture in chondrogenic differentiation medium (CPM). (b) Representative live/dead staining images of the cell-laden constructs at Day 21. Scale bars = (a) 10 mm and (b) 250 µm.

After 21 days of culture, the constructs were harvested for biochemical and histological analyses. Constructs from all groups exhibited showed comparable DNA contents. Furthermore, cartilage-like matrix production was observed in all groups, as evidenced by comparable levels of glycosaminoglycan (GAG) production, an important ECM component of cartilage tissue^48^, and strong whole-construct alcian blue (pH 1.0) staining^49^ (**Figure 9a,b**). Hematoxylin and eosin (H&E) staining revealed relatively uniform cell distribution within the constructs, while Safranin O/Fast Green (SafO/FG) and alcian blue (pH 1.0) staining exhibited strong and comparable GAG deposition across all groups (**Figure 9c**). These results demonstrate that the biaxial gradient-programmed 4D hydrogel system can orchestrate dynamic morphogenesis while simultaneously supporting efficient cartilage-like tissue formation, highlighting its potential for engineering architecturally complex living tissues with programmable developmental trajectories.

**Figure 9.**
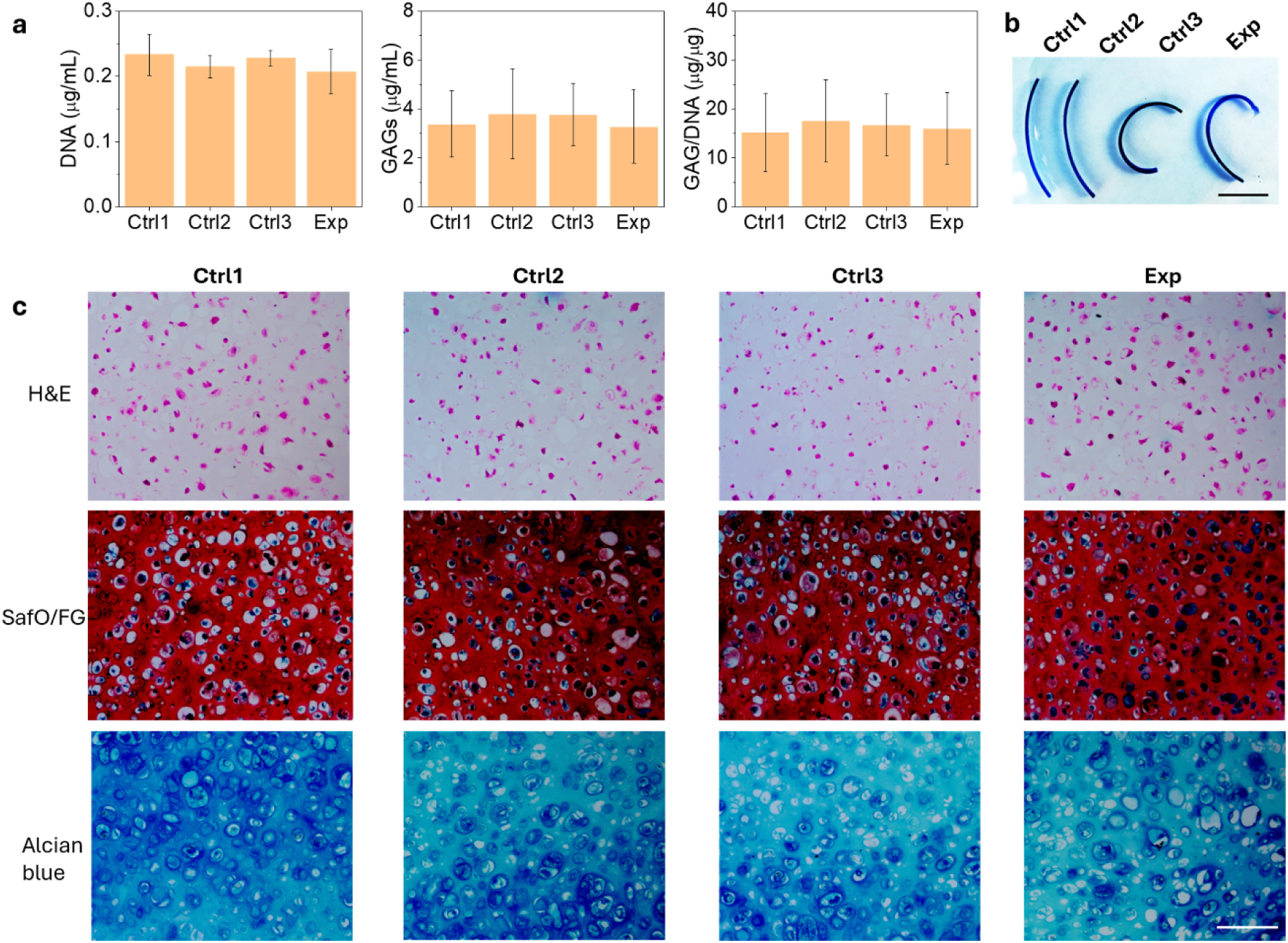
Biochemical and histological characterization of chondrogenically differentiated constructs at Day 21. (a) Quantification of DNA content, GAG production, and GAG/DNA. (b) Whole-construct alcian blue staining (pH 1.0) indicating cartilage-like matrix deposition. (c) Histological staining of sectioned constructs using H&E, SafO/FG, and alcian blue (pH 1.0). Scale bars = (b) 10 mm and (c) 100 µm. Data are presented as mean ± standard deviation (±SD), *N* = 3.

## Conclusion

In summary, a biaxial gradient-programmed 4D hydrogel platform capable of generating spatially coordinated, non-uniform shape morphing within a single construct was developed. Using a one-step photocrosslinking strategy that integrates vertical light attenuation with horizontal grayscale photomask patterning, orthogonal crosslinking gradients were introduced into hydrogels. This design produced spatially heterogeneous swelling stresses, enabling programmable deformation beyond the uniform bending typically observed in conventional uniaxial gradient systems. The biaxial gradient architecture enabled the formation of complex geometries and biomimetic structures, including swan-neck, fiddlehead fern, sea star, and *Euonymus europaeus*-like morphologies. Importantly, the platform supported cell-laden biofabrication, maintaining high cell viability and structural stability during long-term culture. hMSC–laden constructs exhibited efficient cartilage-like ECM deposition under chondrogenic induction. Overall, this work establishes biaxial gradient engineering as an effective strategy for programmable 4D hydrogels, providing a promising platform for morphogenetic tissue engineering and biomimetic biofabrication.

## Materials and Methods

### Polymer synthesis

OMA was synthesized following previously reported protocols with minor modifications^35^. Briefly, to synthesize OMA with a theoretical oxidation of 1% and methacrylation of 30%, alginate (10g, Protanal LF 20/40, FMC Biopolymer) was dissolved in Milli-Q water (900 mL) under stirring overnight at room temperature. A solution containing sodium periodate (NaIO_4_, 0.108 g) in Milli-Q water (100 mL) was rapidly added to the alginate solution and the reaction was allowed to proceed under stirring in the dark at room temperature. Subsequently, 2-morpholinoethanesulfonic acid (MES, 19.52 g, Sigma) and sodium chloride (NaCl, 17.53 g, Sigma) were added, and the pH was adjusted to 6.5 using 5 N sodium hydroxide (NaOH, Sigma). *N*-hydroxysuccinimide (NHS, 1.77 g, Sigma) and 1-ethyl-3-(3-dimethylaminopropyl) carbodiimide hydrochloride (EDC·HCl, 5.84 g, Sigma) were then added and allowed to react for 10 min. Next, 2-aminoethyl methacrylate (AEMA, 2.54 g, Polysciences Inc.) was slowly added and the reaction proceeded overnight in the dark. The product was precipitated in chilled acetone (4 °C), rehydrated with Milli-Q water (100 mL), and dialyzed against Milli-Q water (7 L) for 3 days at 4 °C using dialysis tubing (MWCO 3.5 kDa, Spectrum Laboratories), with the dialysis water changed twice daily. The dialyzed solution was treated with activated charcoal (0.5 mg/100 mL) and filtered through a 0.22 μm membrane. The solution was then frozen at −80 °C overnight and lyophilized for at least 10 days until completely dry. The actual degree of methacrylation (5.6%) was determined by ^1^H NMR spectroscopy (Bruker 600 MHz AVANCE III) using (trimethylsilyl)propionic acid-d_4_ sodium salt (0.05% w/v) as an internal standard in D_2_O (1% w/v), according to previously reported methods.

The GelMA was synthesized following our previously reported protocol^41^ with slight modifications from the original method^50^. Briefly, type-A gelatin (10 g, Sigma) was dissolved in PBS (pH 8.0, 100 mL) at 50 °C under constant stirring until fully dissolved. Methacrylic anhydride (10 mL) was then added dropwise at a rate of 1 mL/min under vigorous stirring. The reaction proceeded for 1 h at 50 °C and was subsequently cooled to room temperature and allowed to react overnight. The crude product was precipitated in excess acetone and purified by dialysis against Milli-Q water using dialysis tubing (MWCO 14 kDa) for 7 days at 50 °C, with the dialysis water changed twice daily. The final GelMA product was obtained after lyophilization for 2 weeks. The actual degree of methacrylation was determined to be 71.8% by ^1^H NMR.

### Photomask Design

Photomasks with defined grayscale values or gradient grayscale ranges were fabricated by printing predesigned grayscale patterns onto transparency films (HP 51636F LX JetSeries) using a standard office inkjet printer (Brother MFC-L5900DW).

### Cell Culture

hMSC were isolated according to previously reported protocols^51^. Cells for non-differentiation experiments were expanded in growth medium consisting of low-glucose Dulbecco’s Modified Eagle Medium (DMEM-LG, Sigma) supplemented with 10% fetal bovine serum (FBS, Sigma) and 1% penicillin/streptomycin (P/S, Gibco). For differentiation experiments, cells were expanded in the same growth medium additionally supplemented with 10 ng/mL fibroblast growth factor-2 (FGF-2, R&D Systems). Cells were cultured in a humidified incubator at 37 °C with 5% CO_2_, and the medium was changed every 2–3 days. Cells at passage 4 (P4) and approximately 80% confluence were used for subsequent experiments.

### Bioink Preparation

OMA/GelMA bioinks were prepared following previously reported procedures^32^ with minor modifications. Briefly, OMA (2% w/v) was dissolved in deionized water and slowly added to a 0.2 M CaCl_2_ solution under vigorous stirring to form ionically crosslinked hydrogel beads.

The beads were collected and washed with 70% ethanol, then mechanically fragmented into microgels using a blender (Osterizer MFG) operating in pulse mode. The resulting microgels were stored in 70% ethanol at 4 °C. Prior to use, the stored microgels were washed three times with Milli-Q water containing photoinitiator (PI, 2-hydroxy-4′-(2-hydroxyethoxy)-2-methylpropiophenone, 0.05% w/v) and PA (4′-hydroxy-3′-methylacetophenone, 0.02% w/v), followed by two washes with DMEM-LG containing the same additives, forming a jammed slurry. GelMA (0.5% w/v) was dissolved directly into the recovered OMA slurry to form the final cell-free bioink. For cell-laden bioinks, hMSCs were mixed into the OMA/GelMA slurry at a density of 10 million cells/mL.

### Rheological Characterization

Rheological properties were measured using a Kinexus Ultra+ rheometer (Malvern Panalytical) with an 8-mm parallel plate geometry and a 1 mm gap. All measurements were conducted at 25 °C. Oscillatory frequency sweep tests (0.1–100 Hz at 1% strain) were performed to determine storage modulus (G′), loss modulus (G″), and viscosity. Oscillatory strain sweep tests (0.01–100% strain at 1 Hz) were conducted to evaluate shear-thinning behavior. Cyclic strain tests alternating between 100% and 1% strain at 1 Hz were performed to assess the recovery of modulus and viscosity (self-healing property).

### Printing

Bioinks were loaded into syringes fitted with 22-gauge needles (McMaster-Carr) and mounted on a BIOX bioprinter (Cellink). Constructs were printed using the parameters listed in **Table S1**. Following printing, constructs were photocrosslinked using UV light (20 mW cm^-2^) for 60 s for large constructs (**Figure 2**) or 30 s for other printed structures with or without photomasks.

Constructs for swelling, morphing, and differentiation experiments were then transferred into culture media and incubated at 37 °C. Cell-free constructs, including disc-shaped samples, were cultured in 24-well plates containing 2 mL medium (H O, PBS, or DMEM-LG) for swelling and mechanical characterization. Strip, star-shaped, and biomimetic 4D architecture constructs were cultured in 6-well plates containing 8 mL PBS and cross-shaped constructs were cultured in 100 mm Petri dishes containing 20 mL PBS for shape-morphing studies. These hydrogels were cultured for 12 h prior to imaging.

### Chondrogenic Differentiation

Hydrogel strips encapsulating hMSCs (10 million cells/mL bioink) were cultured in 6-well plates containing 6 mL chondrogenic medium consisting of high-glucose DMEM (DMEM-HG, Sigma), P/S (1% v/v), ITS^+^ (10% v/v, BD Biosciences), nonessential amino acids (1% w/v, Gibco), sodium pyruvate (100 mM), dexamethasone (0.1 μM), L-ascorbic acid phosphate (129 nM), and transforming growth factor-β1 (TGF-β1, 10 ng/mL, Peprotech). Cultures were maintained for 21 days, with half of the medium replaced daily.

For biochemical analysis, constructs were digested in 0.5 mL papain solution containing papain (50 μg/mL), L-cysteine (2 mM), sodium phosphate (50 mM), and EDTA (2 mM, pH 6.5) at 65 °C for 12 h. Samples were diluted with 0.5 mL nuclease-free water and vortexed on ice.

DNA content was quantified using the PicoGreen dsDNA assay (Invitrogen) with excitation at 480 nm and emission at 520 nm. GAG content was measured using a dimethylmethylene blue (DMMB, Sigma) assay at 595 nm. To account for background signals arising from the OMA/GelMA hydrogel matrix, measurements were calibrated by subtracting the GAG values obtained from acellular OMA/GelMA hydrogel constructs cultured in BPM for 21 days.

Although the acellular constructs do not fully replicate the amount of residual hydrogel present in the corresponding cell-laden constructs after culture, this correction was used as an appropriate approximation to improve the accuracy of GAG quantification. Whole-construct staining was performed by incubating the constructs in alcian blue solution (pH 1.0) for 15 min, followed by immersion in PBS for 1 h at room temperature.

For histological analysis, samples were fixed in neutral buffered formalin (NBF, 10% w/v) overnight at 4 °C, dehydrated, paraffin-embedded, and sectioned at 7 μm thickness using a Leica RM2255 microtome. Sections were stained with H&E, SafO/FG, and alcian blue (pH 1.0). Images were acquired using an Eclipse TE300 microscope (Nikon) equipped with an MU1403 camera (AmScope).

### Statistical Analysis

All experiments were performed using three samples (n = 3) per condition, and data are presented as mean ± standard deviation (SD). Statistical significance was determined using one-way ANOVA followed by Tukey’s HSD post hoc test. Differences were considered statistically significant at *p* < 0.05 unless otherwise specified.

## Author Contributions

A.D.: Conceptualization, Methodology, Investigation, Formal analysis, Investigation, Data curation, Writing - Original Draft, Writing - Review & Editing, Visualization. K.L.G.: Methodology, Investigation,Formal analysis, Writing - Review & Editing, Visualization. E.A.: Conceptualization, Methodology, Formal analysis, Investigation, Resources, Writing - Review & Editing, Supervision, Project administration, and Funding acquisition.

## Competing Interest Statement

The authors declare no competing interest.

## Supporting information

Supporting Inofrmation

## Acknowledgments

The authors gratefully acknowledge funding support from the Department of Veterans Affairs, Veterans Health Administration, Office of Research and Development, Rehabilitation Research and Development Service under award number RX004288. The contents of this publication are solely the responsibility of the authors and do not necessarily represent the official views of the Department of Veterans Affairs.

## Notes

### Competing Interest Statement

The authors have declared no competing interest.

