## Supporting Inofrmation for "4D Biomimetic Morphing Hydrogel Scaffold via Biaxial Gradient Programming"

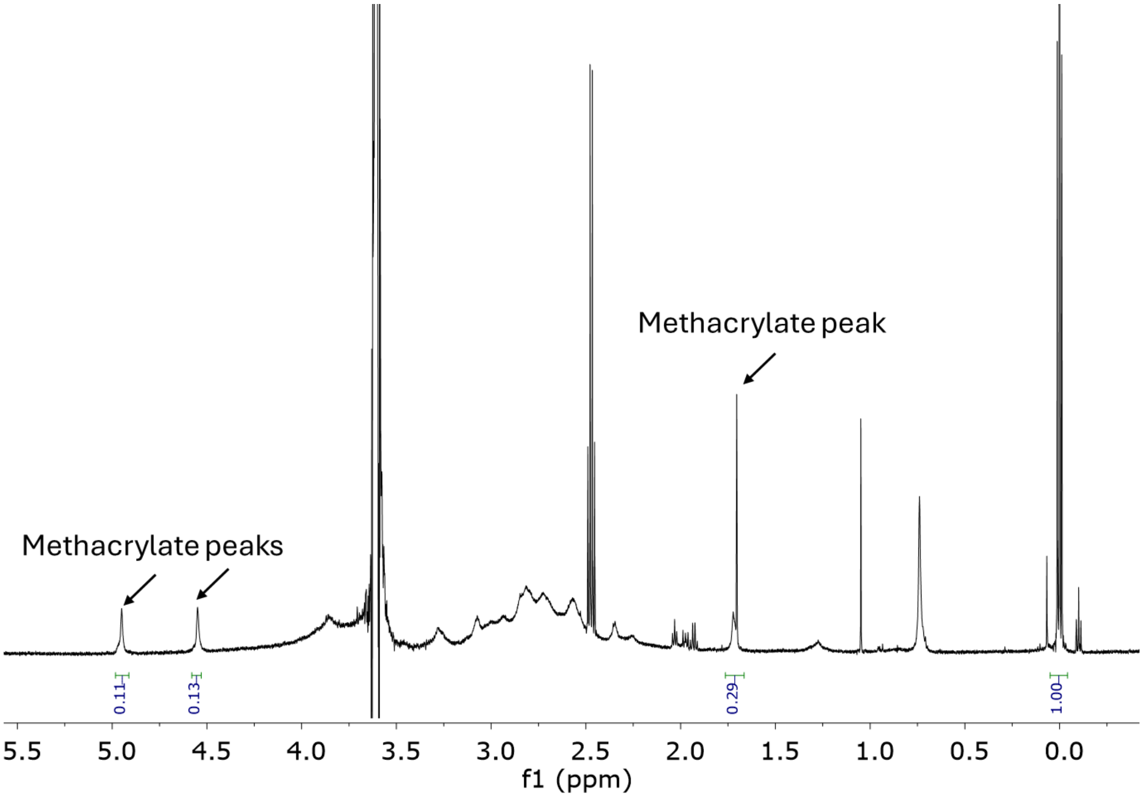


**Figure S1.** The $^{1}$H NMR spectrum of OMA in $D_2O$ (1 w/v %).

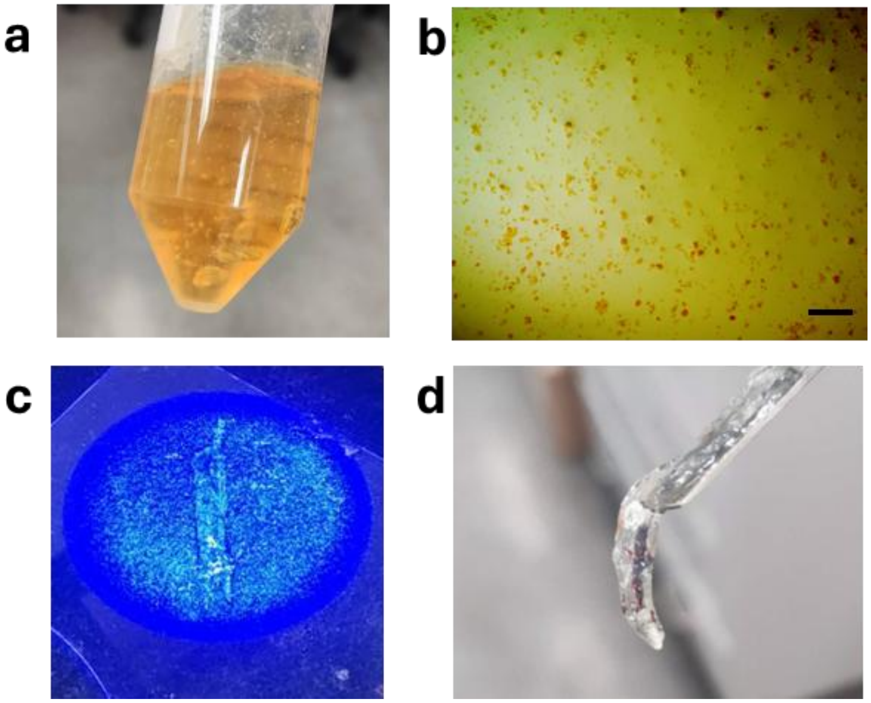


**Figure S2. Jammed slurry microgel bioinks.** (a) As-prepared OMA microgel bioink slurry. (b) Representative morphology of OMA microgels. (c) Photocrosslinking of the OMA microgel bioinks. (d) Photograph of a photocrosslinked OMA hydrogel construct (20 mW $cm^{-2}$, 30 s). Scale bar = 250 µm.

**Table S1**. Printing parameters.

| | |
|---|---|
| **Needle Size** | 22 G |
| **Printing Speed** | 4 mm/s |
| **Extrusion rate** | 1.5 μL/s |
| **Infill Density** | 60% |
| **Layer Height** | 0.66mm |
| **Printing Pattern** | Rectilinear |

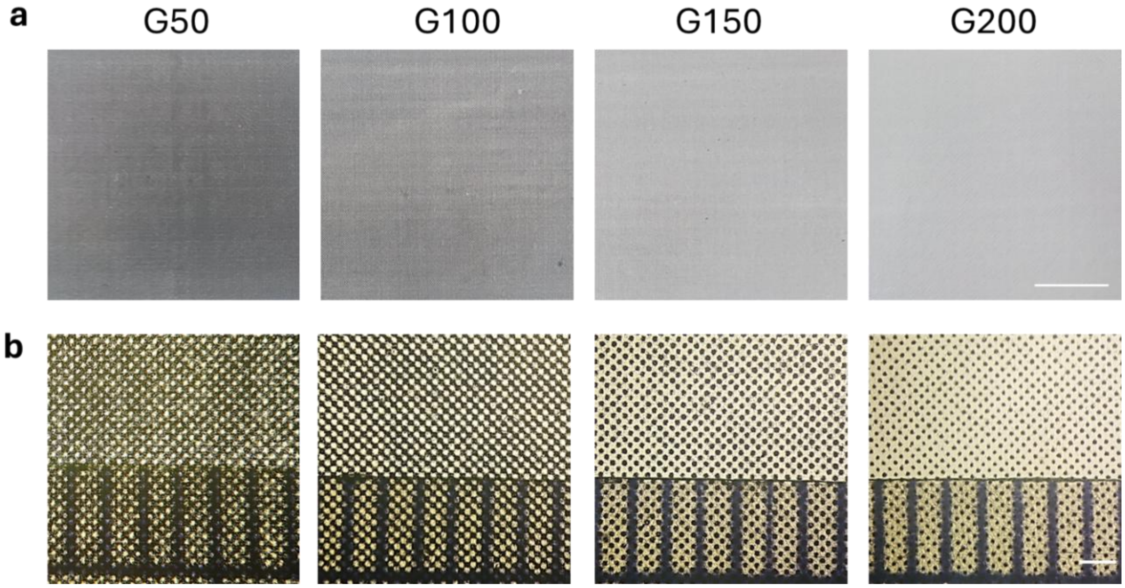


**Figure S3**. Photomicrographs of photomasks with different grayscale values at (a) low and (b) high magnifications. Scale bars = (a) 10 mm and (b) 1.0 mm.

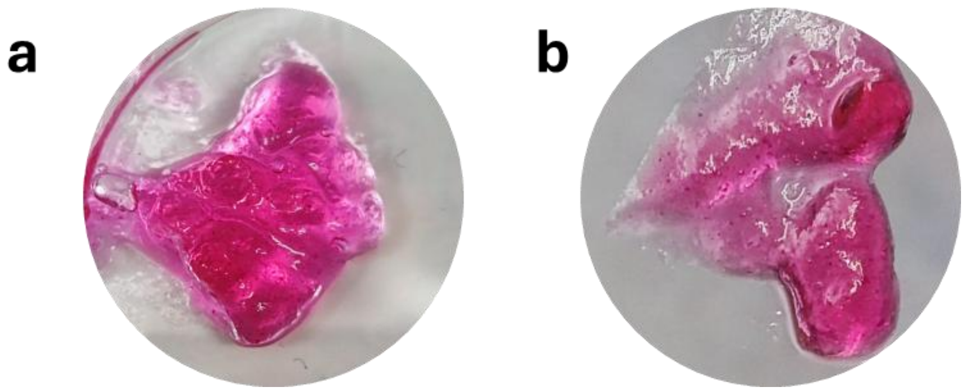

**Figure S4. Photocrosslinking of OMA/GelMA bioinks under different conditions.** (a) Photograph of photocrosslinked bioink prepared using a G50 photomask, showing insufficient structural integrity. (b) Photograph of photocrosslinked bioink prepared under the VG_G150 condition (20 mW $cm^{-2}$, 30 s), showing unstable hydrogel formation. Rhodamine B (0.005% w/v) was incorporated to facilitate visualization.

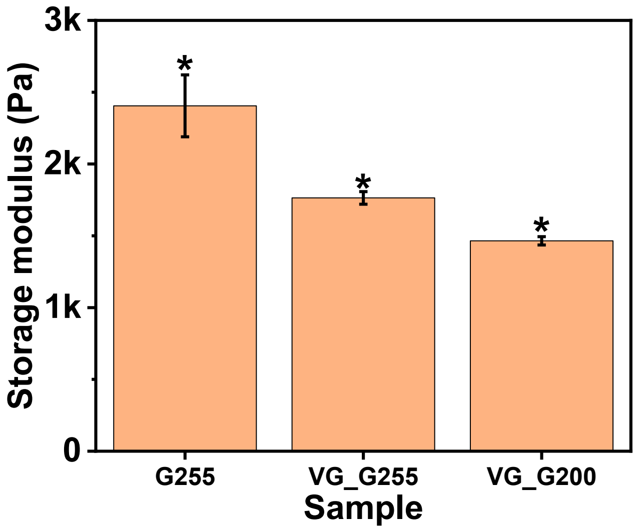


**Figure S5.** Storage modulus (1% strain, 1 Hz) of hydrogels photocrosslinked under different grayscale photomasks with photoabsorber incorporation (VG formation) after swelling in $H_2O$. Data are presented as mean ± standard deviation (±SD), $N = 3$.

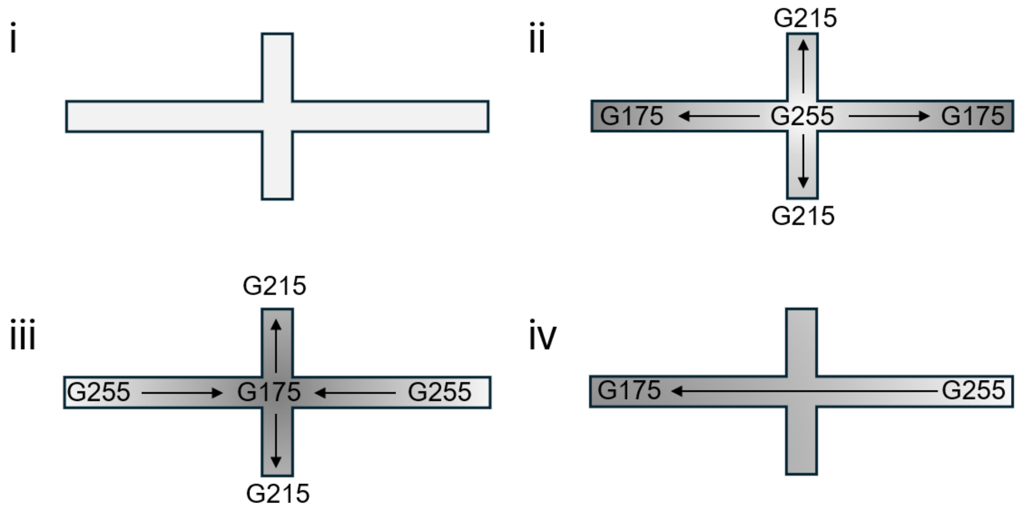


**Figure S6.** HG configuration parameters corresponding to the masks used for Figure 5b.

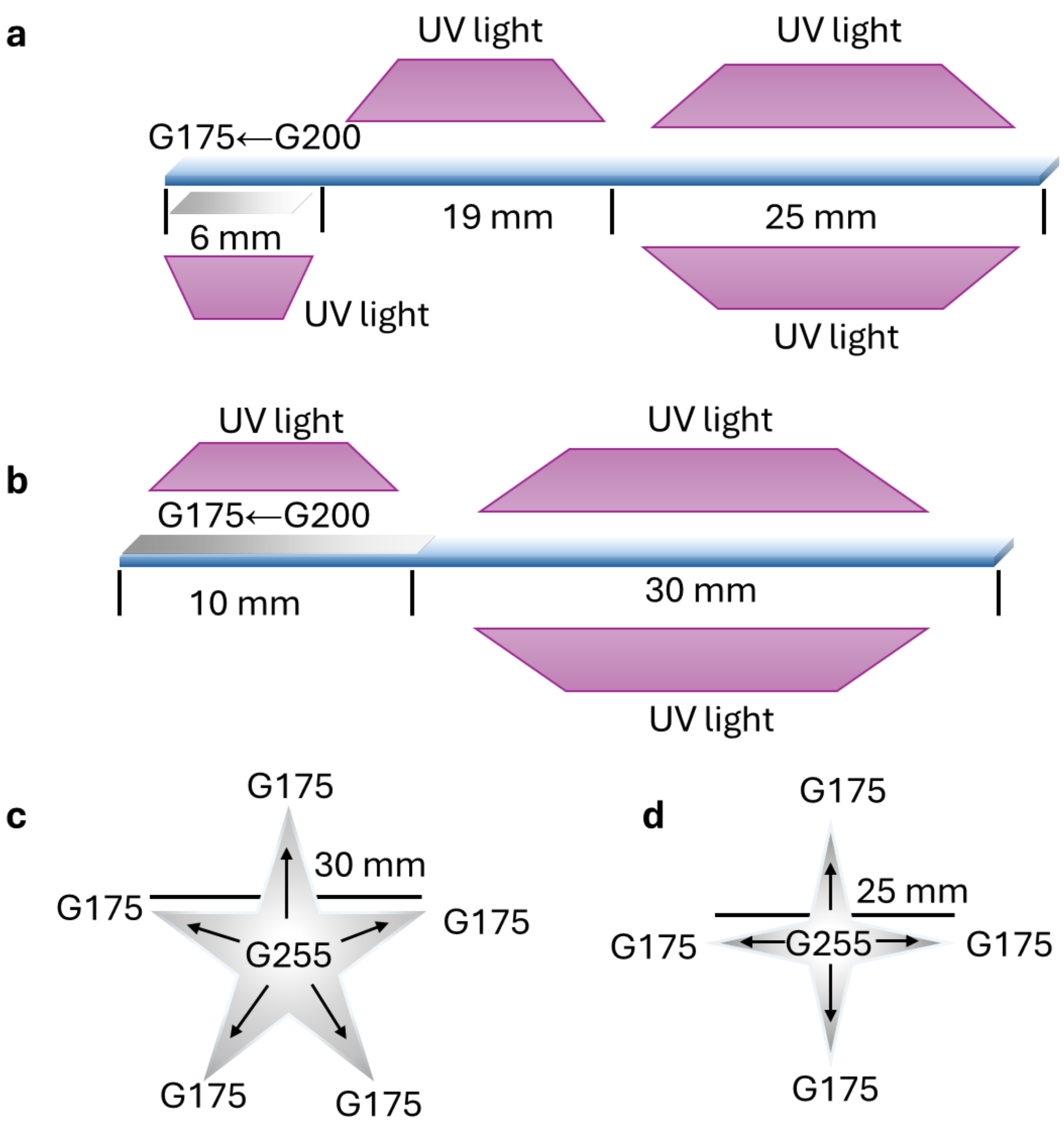


**Figure S7. Design parameters and UV irradiation patterns for the initial structures corresponding to Figure 6.** (a) Swan, (b) fiddlehead fern, (c) five-arm star, and (d) four-arm star.

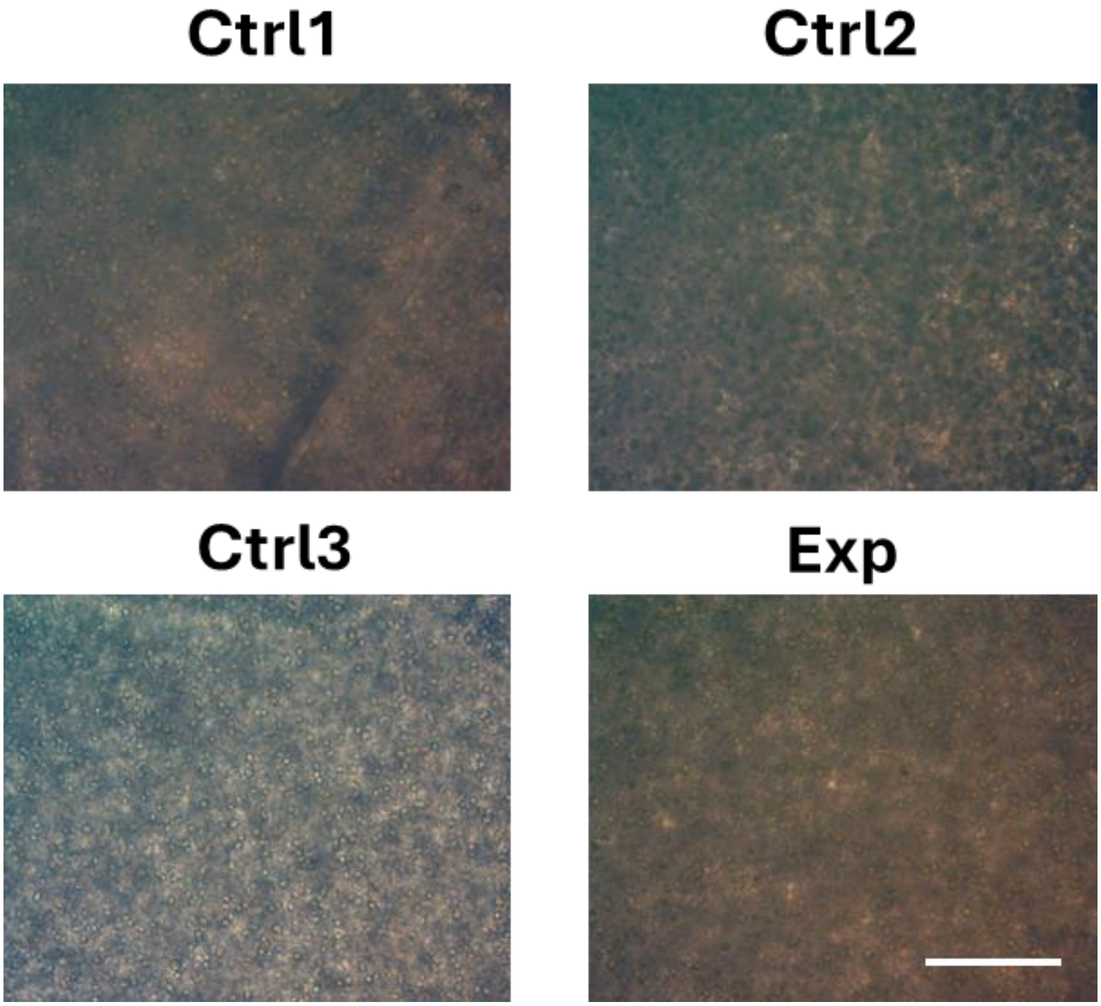


**Figure S8.** Cell morphology within the constructs from different groups at Day 21. Scale bar = 250 μm.
